# Distinct transcriptional programs define a heterogeneous neuronal ensemble for social interaction

**DOI:** 10.1101/2023.12.22.573153

**Authors:** Hailee Walker, Nicholas A. Frost

## Abstract

Reliable representations of information regarding complex behaviors including social interactions require the coordinated activity of heterogeneous cell types within distributed brain regions. Activity in the medial prefrontal cortex is critical in regulating social behavior, but our understanding of the specific cell types which comprise the social ensemble has been limited by available mouse lines and molecular tagging strategies which rely on the expression of a single marker gene. Here we sought to quantify the heterogeneous neuronal populations which are recruited during social interaction in parallel in a non-biased manner and determine how distinct cell types are differentially active during social interactions. We identify distinct populations of prefrontal neurons activated by social interaction by quantification of immediate early gene (IEG) expression in transcriptomically clustered neurons. This approach revealed variability in the recruitment of different excitatory and inhibitory populations within the medial prefrontal cortex. Furthermore, evaluation of the populations of IEGs recruited following social interaction revealed both cell-type and region-specific transcriptional programs, suggesting that reliance on a single molecular marker is insufficient to quantify activation across all cell types. Our findings provide a comprehensive description of cell-type specific transcriptional programs invoked by social interactions and reveal new insights into the heterogeneous neuronal populations which compose the social ensemble.

## Introduction

The medial prefrontal cortex (mPFC) plays a critical role in the regulation of normal social interactions (*1, 2*). Within the mPFC it is the activity of heterogeneous neuronal sub-types which permits the encoding of diverse information relevant to social interactions (*2–4*), anxiety-related behaviors (*5*), and flexible representations of environment (*6*).

Neurons may be segmented by layer, morphological appearance, connectivity, and specific patterns of gene regulation into heterogeneous populations with distinct functional properties within the cortical microcircuit. Normal social interactions require the activity of both excitatory and inhibitory neuron types (*7–9*). The activity of both populations of neurons is critical for normal social behavior (*8, 9*) indicating that social interactions likely utilize computations involving heterogeneous cell types. Single cell datasets have revealed that numerous transcriptionally-defined cell types exist within the laminar cortex (*10*) and are non-randomly connected (*11*) suggesting distinct roles in microcircuit-level computations during behavior. Despite this there has not been a systematic description of the composition of a social ensemble by heterogeneous cell types.

Previous studies quantifying (*4*) or modulating (*8, 9*) the activity of specified populations of neurons during social interaction have provided extraordinary insight into the role of specific molecularly or anatomically defined neurons in computations underlying normal social behavior. However, both electrophysiological and imaging recordings are limited by the number of neuron subtypes which can be assayed simultaneously, as well as the specificity by which neuronal populations can be defined. Similarly, ‘tagging’ approaches (*12, 13*) may be limited in that they typically utilize a single molecular tag to define active populations of neurons which may introduce bias if not all neuron subtypes utilize the same transcriptional program.

We utilized a non-biased transcriptional approach (*14*) to identify heterogeneous neuronal populations within the mPFC and quantify their recruitment during social interactions. Prefrontal neurons isolated from wildtype mice following social interaction showed increased expression of a broad population of immediate early genes (IEGs). Utilizing a panel of socially-responsive IEGs we identified robust and reliable recruitment of activated cells within molecularly-defined glutamatergic and GABAergic populations. Quantification revealed differential neuronal recruitment within distinct excitatory cell types, even within the same cortical layer, highlighting the precision with which neuronal ensembles are recruited during behavior.

## Results

### Transcriptomic Identification of heterogeneous cell types in the mPFC and Cerebellum

Prefrontal microcircuits contain heterogeneous excitatory and inhibitory cell types which play functionally distinct roles in microcircuit function (*11*). We utilized unbiased high-throughput single-nuclei RNA sequencing (snRNA-seq) to transcriptomically identify heterogeneous cell populations of specific brain regions (Figure 1A-C). We isolated nuclei from the mPFC or cerebellum and generated 10X sequencing libraries from which we identified 101,520 cells from the medial prefrontal cortex (mPFC) from 14 adult c57 bl6/J (WT) mice. Identified nuclei were clustered with Seurat and cell types were identified based on known cell-type specific marker genes ((*10, 15*); Figures 1B,C). In total, twenty-two cell types were identified in the mPFC, each with unique transcriptional profiles (Figures 1C,D).

**Figure 1:**
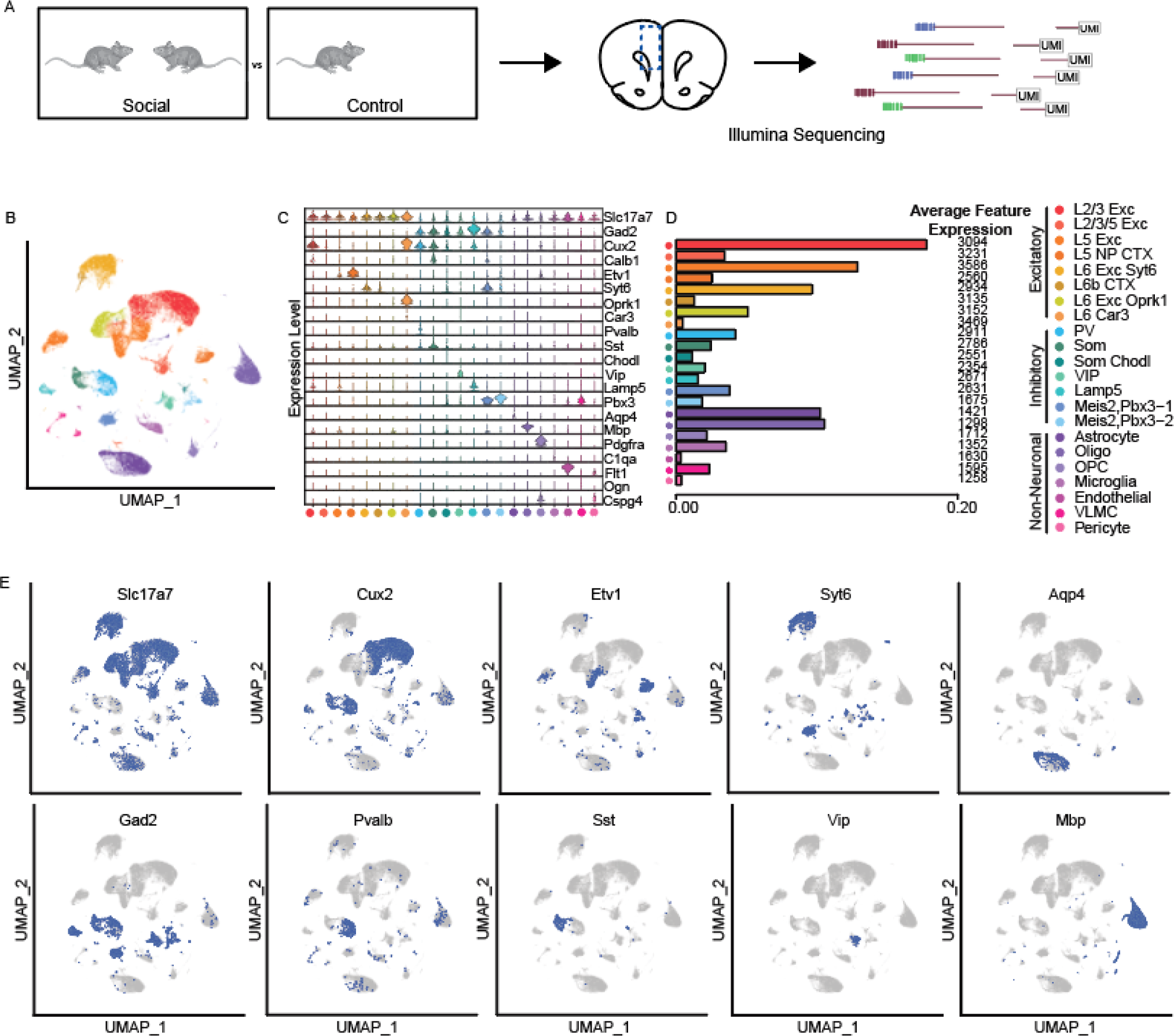
Identification of heterogeneous prefrontal cell types. A) Overview of experimental workflow. B) UMAP visualization of clustered prefrontal cortex cells based on nearest neighbor calculations of 101,520 single-nuclei transcriptomes from WT mice (3 early social, 5 late social; 6 non-social controls). Individual clusters were identified based on gene expression profiles. C) Violin plot showing the expression level of known marker genes across cell types (*10, 15*). D) Bar plot representing the proportion of cells belonging to each cell type and average number of genes expressed by cells of that type (L2/3 Exc – 17.8%, L2/3/5 Exc – 3.5%, L5 Exc – 12.9%, 5 NP CTX – 2.6%, L6 Exc Syt6 – 9.7%, L6b CTX – 1.3%, L6 Exc Oprk1 – 5.1%, L6 Car3 – 0.5%, PV – 4.2%, Som – 2.5%, Som Chodl – 1.1%, VIP 2.1%, Lamp5 - 1.6%, Meis2,Pbx3-1 – 3.8%, Meis2, Pbx3-2 – 1.9%, Astrocytes – 10.2%, Oligodendrocytes – 10.5%, OPC – 2.2%, Microglia 3.6%, Endothelial 0.4%, VLMC – 2.4%, Pericyte – 0.4%). E) UMAP visualization as in B, with cells colored based on high level of expression (> ∼60% max) of given well known, cell-type specific marker genes.

Neurons within the prefrontal cortex are organized into distinct layers (L1, 2/3, 5, and 6) and may be segmented by transcriptomic profile, morphology, and functional properties. We identified excitatory neurons and assigned them to cortical layers in the mPFC using previously identified marker gene profiles (Figure 1E). In total we identified 2 excitatory cell types found in layer 2/3 (L2/3 Exc and L2/3/5 Exc), 2 cell types found in layer 5 (L5 Exc and L5 NP Ctx), and 4 cell types which mapped to layer 6 (or L6 Car3, L6b CTX, L6 Exc Opkr3, and L6 Exc Syt6).

Inhibitory cell type markers Pvalb, Sst, and Vip mapped onto inhibitory clusters marked by expression of Gad2 (Figure 1E). Som Chodl neurons were defined by high expression of Chodl which has been associated with projecting interneurons (*15*). Lamp5 and Meis2,Pbx3 inhibitory neurons express Gad2 and their respective marker genes. Meis2,Pbx3 neurons have previously been identified in the mPFC (*10*) though they are not well-characterized.

Roughly 20% of identified cells were non-neuronal and included Astrocytes (Astro), oligodendrocytes (Oligo), oligodendrocyte progenitor cells (OPC), microglia, endothelial cells, vascular/leptomeningeal cells (VLMC), and pericytes (Figure 1E). Consistent with prior expectations (*10, 16*), excitatory neurons composed 53.24% of all identified cells in the mPFC, while inhibitory neurons made up a smaller (17.1%) proportion.

We identified an additional 90,719 cells from the cerebellum of the same 14 mice (Supp. Figure 1A-C). Cell clusters were identified based on previously described marker genes corresponding to known cerebellar cell types (*17*), which mapped to non-overlapping populations of cerebellar cells (Supp. Figure 1D). We identified seventeen distinct cerebellar cell types in total (Supp. Figure 1A-C). We focused on excitatory and inhibitory neuronal populations in the remainder of this study.

### Identification of region-specific patterns of IEG activation following social interaction

We reasoned that neuronal populations activated during social interaction would display increased immediate early gene (IEG) expression in activated cells as IEGs are quickly and transiently expressed in neurons following neuronal activity and are critical for experience-dependent plasticity (*18*). We pooled cells across mice to examine the expression of a broad panel of IEGs (*14*) in each neuronal cluster in control or social animals and examined the proportion of neurons which expressed at least one copy of each candidate IEG. We considered an IEG to be socially-responsive if the proportion of neurons in at least one cluster increased by at least 2-fold in either social condition (compared to control+1%). We identified a total of 51 socially-responsive IEGs (34 in the mPFC only, 9 in the cerebellum only, 10 in both) (n = 86,559 neurons from 6 control mice, 39,233 neurons from 3 mice 10 minutes following social interaction, and 66,447 neurons from 5 mice 35 minutes following social interaction).

We next examined IEG expression by mouse. Because several clusters contained 0 counts of individual IEGs in one or more condition when examined by mouse we added 1% to both the numerator and denominator. This generated a somewhat larger list than the analysis of pooled data, and we observed an increase in the expression of many IEGs in the social condition compared to control mice (Figure 2B; mPFC: 10-min. 50.3 +/- 0.3 IEGs, 35-min. 45.8 +/- 1.5 IEGs; Cerebellum: 10-min. 31.7 +/- 1.2 IEGs, 35-min. 28.7 +/- 1.5 IEGs) which was significantly higher than that observed in shuffled data (Figure 2B; mPFC: 10-min. 23.0 +/- 2.1 IEGs, 35 min. 23.4 +/- 1.2 IEGs; Cerebellum: 23.3 +/- 1.5 IEGs, 35 min. 22.3 +/- 2.0 IEGs, p <0.0001 for shuffled v. real, 2 way ANOVA). We observed cell-type specific differences in the number of IEGs which increased following social interaction (Figure 2A). Among excitatory neurons, we identified the largest number of socially-responsive IEGs in L6 Car3 and L2/3/5 pyramidal neurons (39 IEGs and 34 IEGs at 10 minutes, respectively) whereas significantly fewer IEGs were identified in L5 NP CTX (20 IEGs at 10 minutes). Within inhibitory populations, LGE-derived Meis2, Pbx3-1 cells contained the most socially-responsive IEGs (33 IEGS increased > 2-fold at 10 minutes). Overall, a larger number of socially-responsive IEGs were identified in the mPFC than in the cerebellum (Figure 2C; mPFC: 10-min. 50.3 +/- 0.3 IEGs, 35-min. 45.8 +/- 1.5 IEGs; Cerebellum: 10-min. 31.7+/- 1.2 IEGs, 35-min. 28.7 +/- 1.5 IEGs, p <0.0001 for region, 2-way ANOVA). This increase in IEG expression was time-dependent such that a greater number of IEGs were positively modulated at 10 minutes post-interaction than at 35 minutes post-interaction, presumably due to egress from the nucleus (Figure 2C; p = 0.0273 for time point, 2-way ANOVA). We restricted further analysis to the more conservative panel of IEGs identified in pooled data.

**Figure 2:**
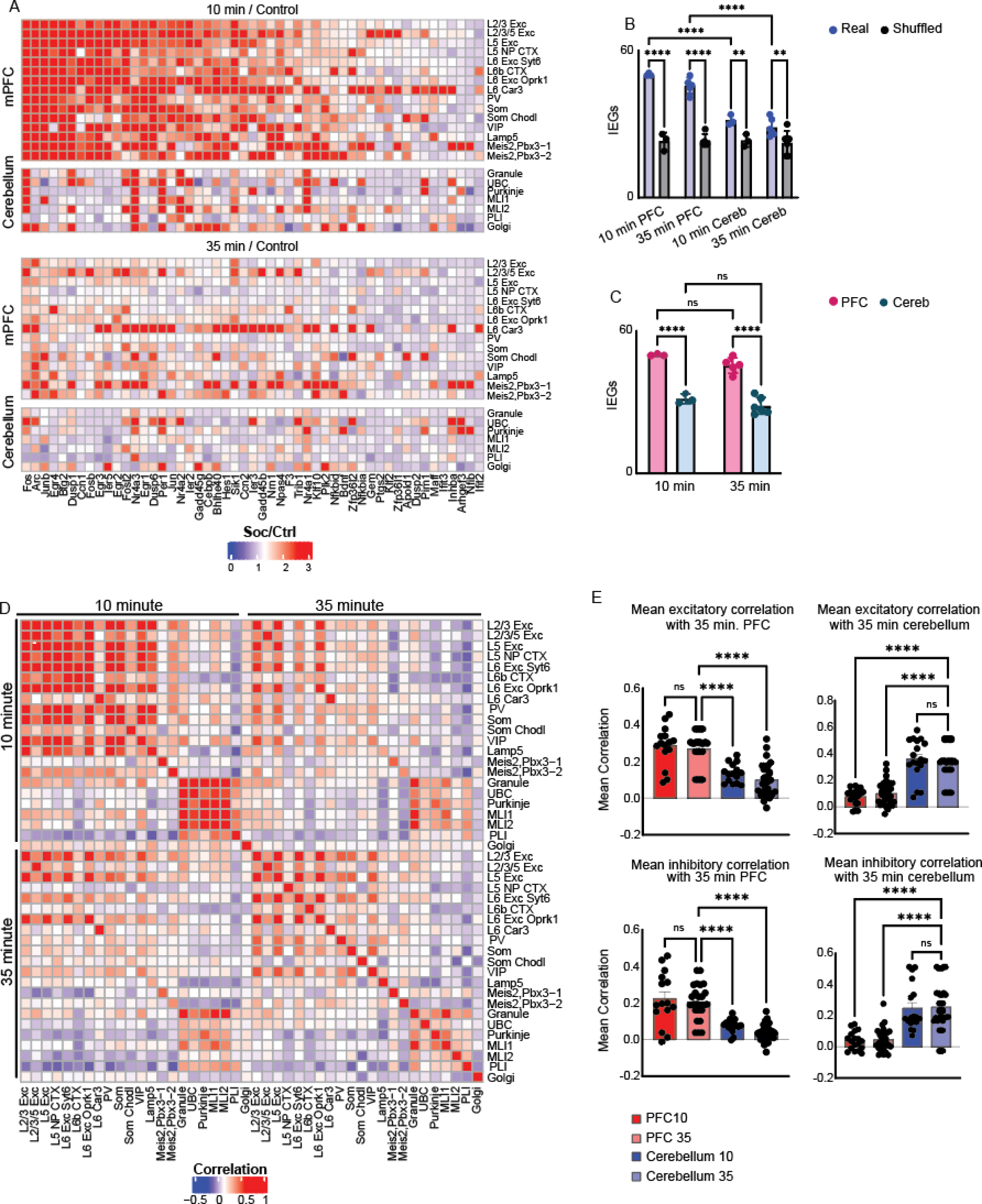
Region-specific patterns of IEG expression following social interaction. We identified 51 IEGs which exhibited greater than 2-fold increase in expression at either 10 or 35 minutes following social interaction (compared to cells from the same cluster in control mice). A) Color plot representing fold-change in the social condition (10 or 35 minute) of each IEG (x-axis) for each cell type within the mPFC or cerebellum. IEGs are sorted based on their fold-increase in mPFC cells. Because some clusters contained 0 counts of a few IEGs in the control condition we added 1% to the denominator for all IEGs. B) Bar graph depicting the number of IEGs per mouse which exhibit 2-fold or greater increase in at least one cell type compared to shuffled data (Real: mPFC 10-min. 50.3 +/- 0.3 IEGs, mPFC 35-min. 45.8 +/- 1.5 IEGs, cerebellum 10-min. 31.7 +/- 1.2 IEGs, cerebellum 35-min. 28.7 +/- 1.5 IEGs; Shuffled: mPFC 10-min. 23.0 +/- 2.1 IEGs, mPFC 35-min. 23.4 +/- 1.2 IEGs, cerebellum 10-min. 23.3 +/- 1.5 IEGs, cerebellum 35-min. 22.3 +/- 2.0 IEGs, p < 0.0001 for real vs shuffled, p < 0.0001 for region and time point, 2-way ANOVA with Sidak’s multiple comparisons). C) Bar graph demonstrating the number of IEGs which exhibit 2-fold or greater increase by region and time (mPFC 10-min. 50.3 +/- 0.3 IEGs, cerebellum 10-min. 31.7 +/- 1.2 IEGs, mPFC 35-min. 45.8 +/- 1.5 IEGs, cerebellum 35-min. 28.7 +/- 1.5 IEGs, p < 0.0001 for region, p = 0.0273 for time point, ANOVA with Uncorrected Fisher’s LSD). D) Correlation matrix showing similarity in IEG expressing proportions for each cell type, region and at each time point. Values were calculated by creating correlation matrix for each social sample/control sample comparison and then averaging by mouse and then across all mice. E) Bar graph of correlation coefficients between samples from each region and time point compared to 35 minute excitatory (top left; 35-minute excitatory mPFC vs: 10-minute mPFC, 0.29 +/- 0.03 correlation coefficient, p = 0.903, 35-minute mPFC, 0.27 +/- 0.02 correlation coefficient, 10-minute cerebellum, 0.14 +/- 0.01 correlation coefficient, p < 0.0001, 35-minute cerebellum, 0.10 +/- 0.02 correlation coefficient, p < 0.0001, one-way ANOVA with Sidak’s multiple comparisons) and inhibitory mPFC neuron populations (bottom left; 35-minute inhibitory mPFC vs: 10-minute mPFC, 0.22 +/- 0.04 correlation coefficient, p = 0.890 , 35-minute mPFC, 0.20 +/- 0.02 correlation coefficient, 10-minute cerebellum, 0.08 +/- 0.01 correlation coefficient, p < 0.0001, 35-minute cerebellum, 0.04 +/- 0.01, p < 0.0001, one-way ANOVA with Sidak’s multiple comparisons) or 35-minute excitatory (top right; 35-minute excitatory cerebellum vs: 10-minute mPFC, 0.08 +/- 0.01 correlation coefficient, p < 0.0001, 35-minute mPFC, 0.10 +/- 0.02 correlation coefficient, p < 0.0001, 10-minute cerebellum, 0.36 +/- 0.03 correlation coefficient, p = 0.980, 35-minute cerebellum, 0.35 +/- 0.02 correlation coefficient, one-way ANOVA with Sidak’s multiple comparisons) and inhibitory cerebellum neuron populations (bottom right; 35-minute inhibitory cerebellum vs: 10-minute mPFC, 0.04 +/- 0.01 correlation coefficient, p < 0.0001, 35-minute mPFC, 0.04 +/- 0.01 correlation coefficient, p < 0.0001, 10-minute cerebellum, 0.25 +/- 0.03 correlation coefficient, p = 0.996, 35-minute cerebellum, 0.26 +/- 0.03 correlation coefficient, one-way ANOVA with Sidak’s multiple comparisons) with standard error.

We next examined the degree of similarity between the population of IEGs which were socially-responsive across different cell types and conditions (Figure 2D,E). Pooled excitatory neurons in the mPFC displayed similar changes in IEG expression at 10 and 35 minutes which diverged from changes in expression observed in the cerebellum. (Excitatory mPFC neurons: 35-min mPFC vs. 10-min mPFC, correlation coefficient 0.29 +/- 0.03, 35 minute mPFC vs 35 minute mPFC correlation coefficient 0.27 +/- 0.02, 35 minute mPFC vs 10 minute cerebellum correlation coefficient 0.14 +/- 0.01, 35 minute mPFC vs 35 minute cerebellum correlation coefficient 0.10 +/- 0.02, p < 0.0001, ANOVA). Similarly, we observed a greater similarity between IEG expression in pooled cerebellar excitatory neurons from the cerebellum at 10 minutes and cerebellar populations at 35 minutes than to mPFC expression at either timepoint (Excitatory cerebellar neurons: 35-min cerebellum vs. 10-min cerebellum correlation coefficient 0.36 +/- 0.03, 35 minute cerebellum vs 35 minute cerebellum correlation coefficient 0.35 +/- 0.02, 35 minute cerebellum vs 10 minute mPFC correlation coefficient 0.08 +/- 0.01, 35 minute cerebellum vs 35 minute mPFC correlation coefficient 0.10 +/- 0.02, p < 0.0001, ANOVA). We observed similar findings when we examined inhibitory populations in the mPFC and the cerebellum at each timepoint (Figure 2D,E). This suggests that region-specific patterns of IEG expression are more distinct than cell-specific patterns of IEG expression.

To further examine the similarity in IEG expression between broad neuronal cell type (excitatory and inhibitory) in the mPFC and cerebellum, we performed principal component analysis (PCA) on the expression of 51 socially-responsive IEGs. The first three principal components accounted for 80.0% of the variance, with differences between the cerebellum and mPFC, as well as between excitatory and inhibitory neuron types, accounting for the majority of IEG expression differences (Figure 3A). Examination of vectors drawn from the mean of the control condition to the mean of either social group suggested divergent patterns of IEG activation occurred in mPFC and cerebellar neurons (Figure 3B,C).

**Figure 3:**
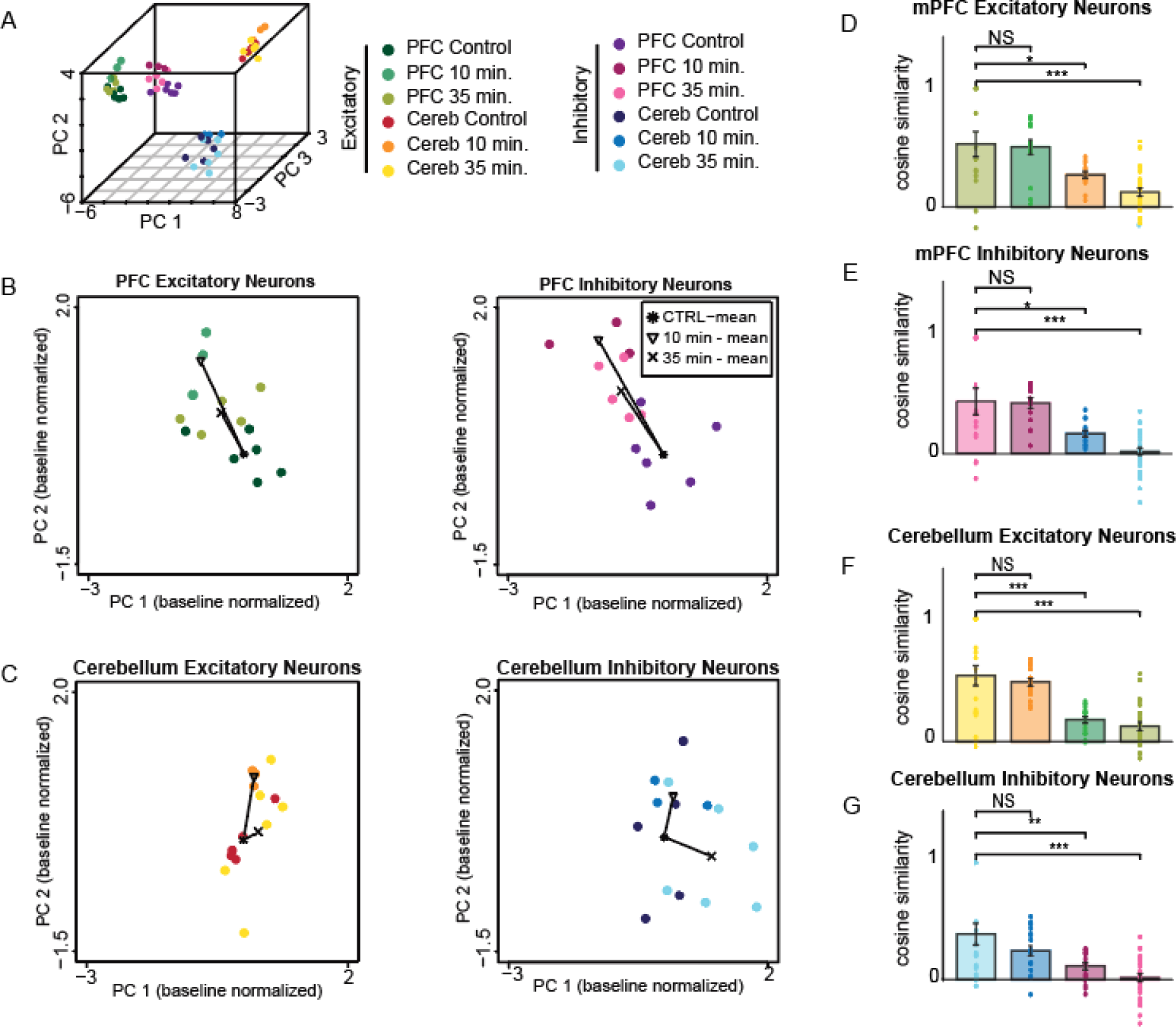
Comparison of region-specific and cell-type specific patterns of IEG expression following social interaction. A) IEG expression averaged across excitatory or inhibitory neuron types for each region and time point plotted following principal components analysis. Each point represents a sample from an individual mouse plotted in principle component space revealing regional and cell type (excitatory vs inhibitory) specific differences in baseline IEG expression. B-C) All points were normalized to the mean of the control condition. B and C demonstrate normalized values for each mouse colored by control and social conditions. Both the mean position of each condition, and the trajectory between the mean control and social conditions are overlaid, demonstrating similar trajectories for each mPFC timepoint and cell-type, which are distinct from trajectories observed in the cerebellum for excitatory or inhibitory neurons. D-G) Bar plots depicting cosine similarity for each social condition compared to all other conditions within the same neuronal subtype for excitatory mPFC (D; 35 min excitatory mPFC similarity with: 35-min. mPFC, 0.53 +/- 0.11, 10-min. mPFC, 0.51 +/- 0.06, 10-min. cerebellum, 0.27 +/- 0.03, 35-min. cerebellum, 0.13 +/- 0.03, p < 0.0001, ANOVA with Dunnett’s multiple comparisons test), inhibitory mPFC (E; 35-min inhibitory mPFC similarity with: 35-min. mPFC, 0.45 +/- 0.12, 10-min. mPFC, 0.43 +/- 0.05, 10-min. cerebellum, 0.17 +/- 0.05, 35-min. cerebellum, 0.02 +/- 0.03, p < 0.0001, ANOVA with Dunnett’s multiple comparisons test), excitatory cerebellum (F; 35 min excitatory cerebellum similarity with: 35-min. cerebellum, 0.54 +/- 0.08, 10-min. cerebellum, 0.48 +/- 0.03, 10-min. mPFC, 0.18 +/- 0.03, 35-min. mPFC, 0.13 +/- 0.03, p < 0.0001, ANOVA with Dunnett’s multiple comparisons test), or inhibitory cerebellum (G; 35 min inhibitory cerebellum similarity with: 35-min. cerebellum, 0.39 +/- 0.09, 10-min. cerebellum, 0.25 +/- 0.04, 10-min. mPFC, 0.11 +/- 0.03, 35-min. cerebellum, 0.02 +/- 0.03, p < 0.0001, ANOVA with Dunnett’s multiple comparisons test).

To quantify these similar and divergent patterns of IEG expression we calculated the cosine similarity between vectors drawn between individual control samples, and social samples from either group (10-minute or 35-minute; Figure 3D-G). As before this revealed a greater similarity between IEG expression in pooled excitatory neurons within the mPFC at 10 minutes and IEG expression within the mPFC at 35 minutes than between the mPFC and either cerebellar timepoint (Excitatory mPFC neurons: 35-min mPFC vs. 10-min mPFC, cosine similarity 0.51 +/- 0.06, 35 minute mPFC vs 35 minute mPFC cosine similarity 0.53 +/- 0.11, 10 minute cerebellum vs 35 minute mPFC cosine similarity 0.27 +/- 0.03, 35 minute cerebellum vs 35 minute mPFC cosine similarity 0.13 +/- 0.03, p < 0.0001, ANOVA). Similarly, we observed a greater similarity between IEG expression observed in the cerebellum at 10 minutes and 35 minutes than between the cerebellum and the mPFC at either time point (Excitatory cerebellar neurons: 35-min cerebellum vs. 10-min cerebellum correlation coefficient 0.48 +/- 0.03, 35 minute cerebellum vs 35 minute cerebellum correlation coefficient 0.54 +/- 0.08, 35 minute cerebellum vs 10 minute mPFC correlation coefficient 0.18 +/- 0.03, 35 minute cerebellum vs 35 minute mPFC correlation coefficient 0.13 +/- 0.03, p < 0.0001, ANOVA). We observed similar findings within inhibitory populations in the mPFC and the cerebellum across timepoints (Figure 3D-G). Taken together these data suggest that regional differences in IEG-driven transcriptional programs exceed cell-type specific differences within a given region.

### The heterogenous social ensemble

We next aimed to quantify the overall change in the active proportion of each cell type following social interactions (Figure 4 & Supp. Figure 2). Individual IEGs were identified in a small proportion of cells (roughly 10%) following social interaction. Moreover, the degree to which individual IEGs were upregulated was not uniform across cell types. For instance, whereas increased Arc expression was observed in the majority of mPFC excitatory neurons, its expression only increased in 2 cerebellar cell types (Figure 4 & Supp. Figure 2). This suggests that identification of activated cell populations using only a single marker gene may be insufficient to compare activation between different types of neurons.

**Figure 4:**
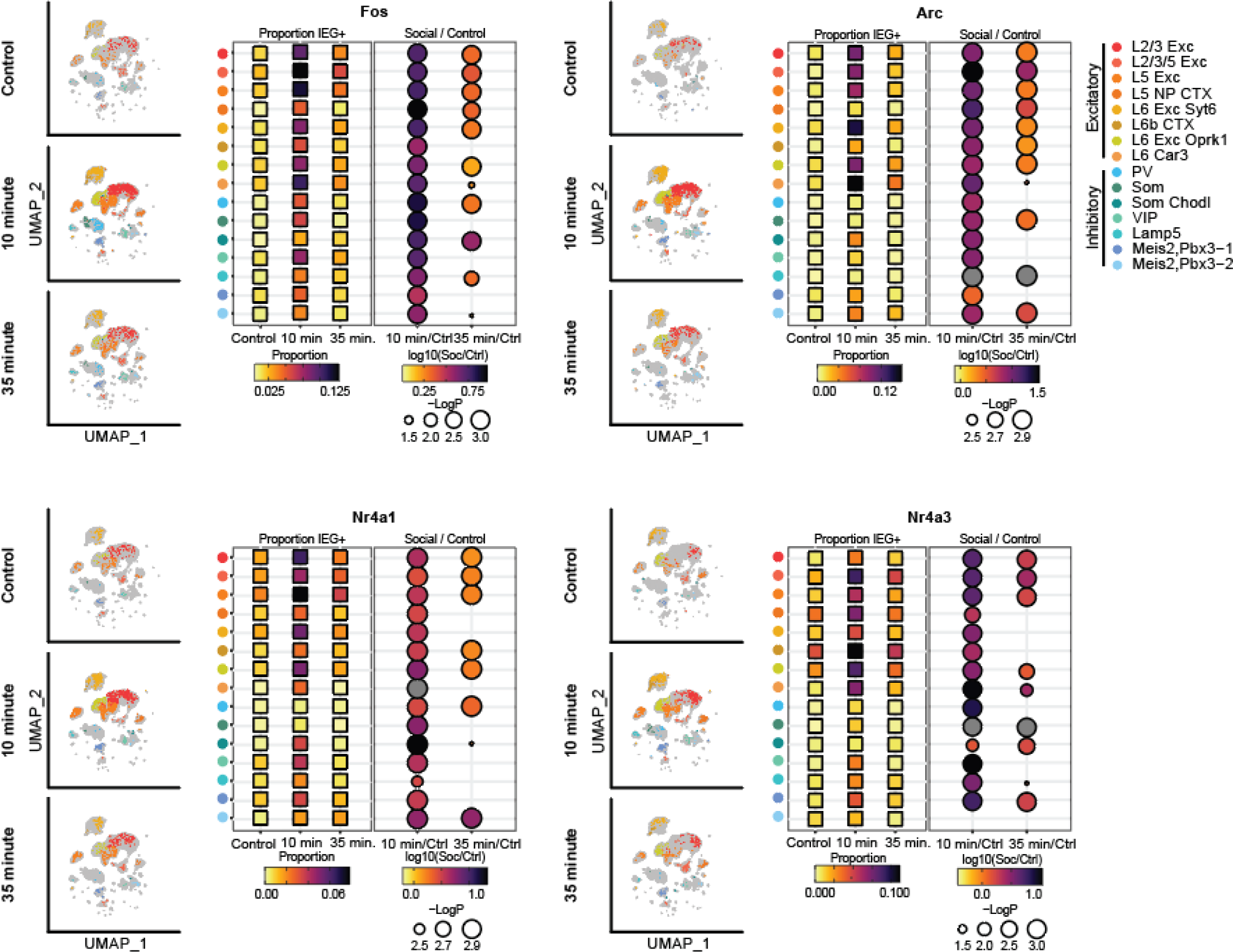
IEG expression in heterogeneous prefrontal neuron populations following social interaction. Cells expressing Fos, Arc, Nr4a1, and Nr4a3 (> 95^th^ percentile of control) are plotted for each timepoint and quantified. Individual colored dots represent individual neurons expressing corresponding IEG, overlaid on clustered dataset randomly down sampled to show equivalent cell counts in each condition. Corresponding dot plots represent the proportion of cells expressing the corresponding IEG at 10 minutes and 35 minutes post social interaction, and the ratio of the proportions of positive cells in the post-social vs control conditions (also see Supp. Table 1; p-values calculated based on shuffled data).

We therefore reasoned that activated cells (IEG+ cells) within different clusters could be more reliably detected using a region-specific panel of IEGs than choosing a single gene. We utilized all IEGs identified as socially-responsive within a given region (defined as > 2-fold change in pooled data). This corresponded to a total of 43 IEGs in the mPFC and 18 IEGs in the cerebellum (Figure 5A; Supp. Figure 3A). Some socially-responsive IEGs were identified in both cerebellar and mPFC cell-types (Arc, Dusp1, Fos, Ler2, Junb, Nr4a1-3, Per1), others (Klf2, Nfknbid, Pim1, Plk2, Sik2, Arhgef3, Inhba, Nfib, Supp. Figure 3) increased significantly only in cerebellar neurons, and still more were only identified in mPFC datasets (Apold, Bhlhe40, Btg2, Ccn1, Ccn2, Cebpb, Dusp2, Dusp6, Egr1-4, F3, Fosb, Fosl2, Gadd45b, Gadd45g, Gem, Hes1, Ler3, Ler5, lfit2, lfit3, Jun, Klf10, Maff, Nfkbia, Npas4, Nrn1, Ptgs2, Sik1, Trib1, Zfp36l1&2; Figure 5).

**Figure 5:**
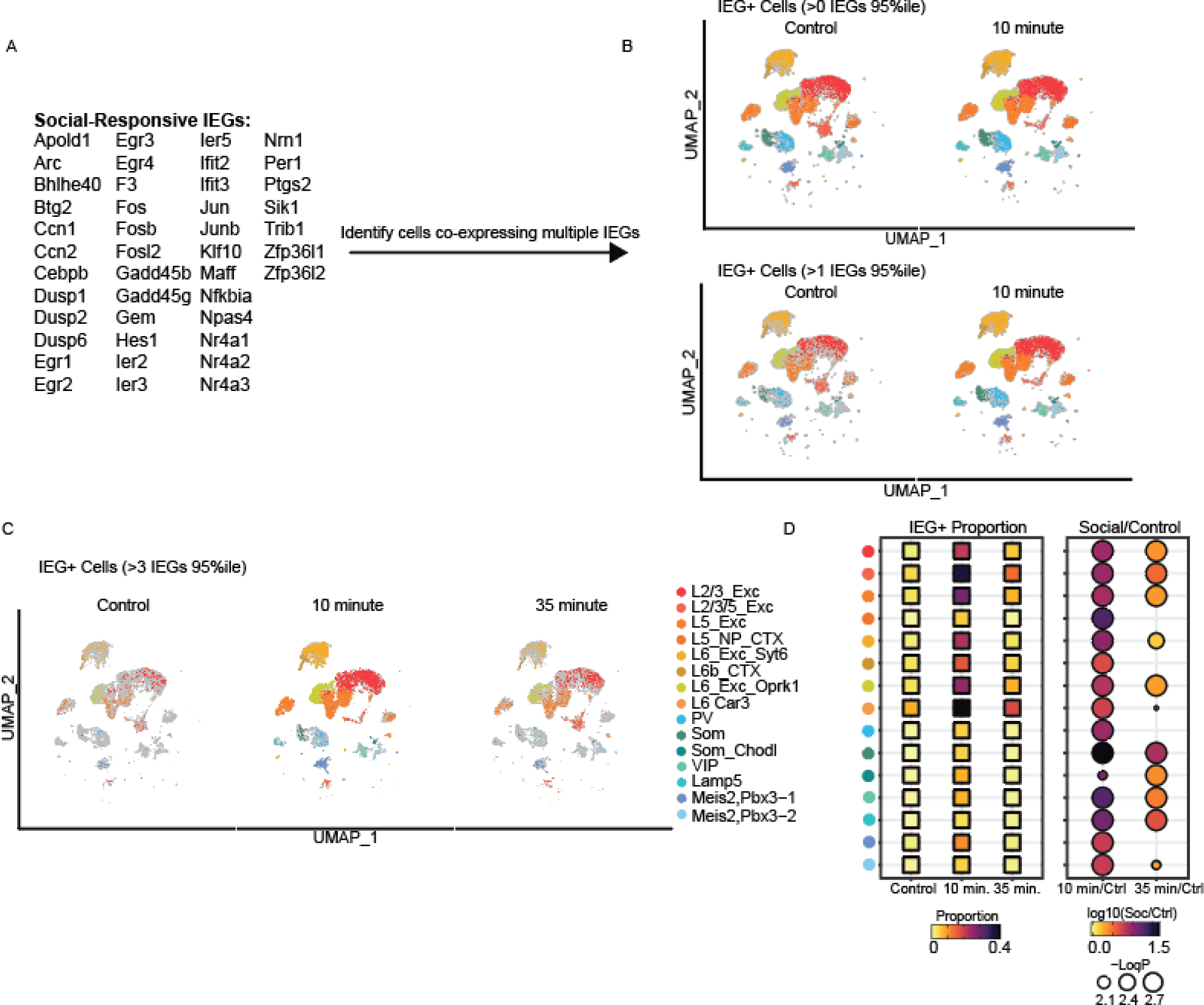
IEG expression defines a heterogeneous social ensemble in the prefrontal cortex. A) We utilized a panel of 43 socially-responsive IEGs in the mPFC (>2-fold expression in social condition compared to control). B) UMAPs showing the proportion of cells of each type which express at least a single IEG (top) or 2 or more IEGs (bottom) in the control and 10-minute post-social condition. C) Proportion of cells expressing 4 or more IEGs above the 95th percentile of expression in the control group. UMAPs are randomly down sampled by condition to show the same number of cells for each condition (B&C). D) Boxes on left are colored to represent the proportion of IEG+ neurons in each cell type in control or social conditions (n = 31,290 neurons from 6 control mice, 24,036 neurons from 5 35-minute post social mice and 16,121 neurons from 3 10-minute post social mice). We then calculated the proportion of IEG+ neurons at each post-social timepoint. Circles are colored to represent the ratio of IEG+ cells in each social condition relative to the control condition. The diameter of each circle represents Log10 p-value (also see Supp. Table 2; p-values calculated based on shuffled data).

IEG+ cells in the mPFC were defined by high expression of 4 or more socially-responsive IEGs (> 95^th^ percentile of control expression; Figure 5B,C). We then calculated the proportion of IEG+ cells in each cluster to quantify heterogeneous cell populations activated by social interaction. We observed a significant increase in the proportion of IEG+ neurons in excitatory and inhibitory prefrontal cell clusters at both 10 and 35 minutes following social interaction (compared both to control and shuffled data; Figure 5D).

We defined IEG+ neurons in the cerebellum as those having high expression of 2 or more socially-responsive IEGs (> 95^th^ percentile of control expression; Supp. Figure 3B,C). The proportion of IEG+ cells increased in both excitatory and inhibitory neurons at 10 and 35 minutes following social interaction (compared to control or shuffled data), but the increase was blunted compared to that observed in the mPFC (10 min mPFC, 12.6 +/- 2.8 fold-increase, 10 min Cerebellum, 3.6 +/- 0.5 fold-increase, 35-min mPFC 2.5 +/- 0.5 fold-increase, 35-min Cerebellum 1.6 +/- 0.3 fold-increase, p = 0.04 for region & p = 0.013 for time point, 2-way ANOVA), potentially reflecting restricted or regional recruitment of cerebellar activity during social interaction.

The relative abundance and the proportional increase in IEG+ neurons varied by cell type within both the mPFC and cerebellum. Even within a given layer, we observed unequal recruitment of different cell types (for instance L5 Exc vs L5 NP CTX, or L6 Car3 vs L6b CTX; Figure 5D). Examination of the total number of cells of each type which composed the social ensemble also revealed the importance of cell abundance; cerebellar granule cells - due to their abundance - were the largest component of the social ensemble (19.3% of significantly active cells 10-minutes post social interaction, and 42.1% of significantly active cells 35-minutes post social interaction; Supp. Figure 4).

## Discussion

Normal social interactions depend on reliable encoding of social information by the prefrontal cortex. During social interactions, information relevant to these interactions must be received, decoded, and routed through the mPFC to distributed brain regions. Thus, the neuronal populations which we define as neuronal ensembles consist of heterogeneous cell types which play distinct roles in the processing and communication (*4, 19*) of information within brain circuits. Cells within the prefrontal microcircuit are nonrandomly connected (*11*), and perturbing the recruitment of one cell type may have unexpected consequences on network function. Indeed, altered recruitment of even a single cell type has been demonstrated to alter social behavior (*8, 9*), presumably by disrupting computations within the prefrontal microcircuit.

This study builds on prior studies which have collectively demonstrated critical features of ensemble representations. First, ensembles are composed of neurons which may be reliably activated or repressed during social interaction (*2, 3, 6*), permitting the generation of social ensembles which are orthogonal to representations of other types of information despite utilizing overlapping cells. This oppositional activation and repression of activity within the mPFC is most likely driven at least in part by recurrent inhibition. While prior studies have implicated specific inhibitory neuronal populations in the regulation of social interactions (*7, 8, 20, 21*) our data supports a model in which heterogeneous inhibitory neuron populations are recruited alongside excitatory neurons during social interaction.

Second, selective routing of activity by defined projections during complex behaviors including social interactions provides an anatomic substrate for distributed processing of social information as well as the top-down control of social interactions by prefrontal circuits (*22, 23*). We observe activation of pyramidal neurons within all cortical layers during social interaction, arguing that anatomic segregation of particular projections (*4*) is insufficient to achieve this selective recruitment which may instead be accomplished through distinct patterns of inhibitory (*11, 24–26*) and excitatory input (*11*).

Critically, our data emphasize complex and heterogeneous transcriptional programs are evoked within single neurons. In our dataset, we observed recruitment of multiple IEGs within individual neurons following social interaction, and the population of socially-responsive IEGs varied markedly between the prefrontal cortex and cerebellum. Even within regions, individual IEGs varied in their ability to identify heterogeneous populations of neurons. For example, whereas Fos expression increased across both excitatory and inhibitory neurons, Arc expression increased less reliably in inhibitory neuron populations.

The source and types of information being processed within the mPFC during social interactions are critical but not fully understood, and includes information related to sensory stimuli (*27*) and internally generated states. These latter states may include information related to recollection or prior experience (*28*), social hierarchy (*29*) or competing motivations (*30*). Critically, different inputs may differentially activate distinct populations of prefrontal neurons resulting in the recruitment of distinct computational units or cell populations (*31*). Our data suggest that these computations utilize distinct excitatory and inhibitory populations to different degrees, which may be critical for generating distinct representations of distinct types of behavioral information.

## Methods

### Animals

Nuclei suspensions from mouse prefrontal cortex (mPFC) and cerebellum were collected from 14 C57BL/6J male and female mice aged 4-5 months. Mice were group housed in our vivarium on a 12- hour light dark cycle with ad libitum food and water until isolation 3 days prior to tissue collection. Novel juveniles (C57BL/6J) used in social interaction assay were sex-matched and aged 4-6 weeks. All animal use was approved by the University of Utah Institutional Animal Care and Use Committee.

### Social Interaction Assay

Three days prior to the day of sequencing mice were singly housed to reduce social-interaction related neural activity prior to the behavioral assay. In addition, one day prior all mice were habituated to the opening of their cage lid by lifting each lid for about 10 seconds and then replacing it every 5 minutes for two hours. One hour prior to euthanasia, half of the cohort were subject to a social interaction assay, while control mice were subjected to sequential cage lid removal to match lid openings experienced by the social cohort. Social mice engaged in sequential 5-minute blocks of social interactions with 5 novel sex-matched juveniles. At a specified time point following removal of the final mouse (or final lid opening) mice were euthanized by isoflurane overdose immediately followed by decapitation and tissue dissection. To prevent light-cycle induced changes in activity or IEG expression all behavioral experiments were performed between 6:00 – 7:00 AM. Tissue was therefore harvested between 7:00 – 8:00 AM for all animals.

### Single-nuclei RNA sequencing

Mice were deeply anesthetized by administration of isoflurane in a gas chamber flowing at 5% isoflurane. Anesthesia was confirmed by negative toe pinch response. Following decapitation whole brains were removed and quickly submerged in ice cold 1X phosphate buffered saline with 15 uM actinomycin-D to prevent artificially induced transcriptional perturbation (*14*). Nuclei were isolated using a protocol developed based on previously described methods (*32*). Briefly, regions of interest were micro-dissected and transferred into 15 uM actinomycin-D (Sigma-Aldrich #A1410-10MG) containing lysis buffer and homogenized with a pestle. Samples were spun down and supernatant was removed and replaced with new lysis buffer with Triton-X (MilliporeSigma #NUC201) two additional times. Samples were filtered through a 40 uM Flowmi cell strainer (Fisher Scientific #14100150). Nuclei are isolated and myelin was removed using a sucrose gradient (MilliporeSigma #NUC201). Nuclei were then resuspended in a resuspension buffer containing 0.5% Ultrapure BSA (Fisher Scientific #AM2616) to prevent clumping of nuclei and 0.2 U/ul RNAse inhibitor (New England Biolabs #M0314S) to prevent RNA degradation. Samples were spun down, supernatant removed and resuspended in 100 uL resuspension buffer, and filtered a final time through a 40 uM cell strainer. Nuclei counts were calculated on a Countess3 and Nexcellom cell counter and samples were diluted to around 700-1,000 nuclei/uL. All samples remained on ice throughout this process when outside of the centrifuge. Samples were delivered to the University of Utah High-Throughput sequencing core immediately following nuclei isolation for library preparation.

Barcoded snRNA cDNA libraries were generated for each sample with the 10x Genomics Chromium Next GEM Single Cell 3’ Gene Expression Library prep v3.1, targeting 7,000-10,000 cells per sample. Libraries were generated following the 10X protocol, with the exception of extending the PCR elongation step from 1 to 3 minutes (*33*). The prepared cDNA library was then sequenced on the Illumina NovaSeq 6000. We generated 200-300 M reads per sample and mapped them to the GRCm39 with the 10x Genomics Cell Ranger software (v4.0.8).

### Cell type clustering

Filtered gene expression matrices generated by Cell Ranger were clustered using Seurat (v4.2.1) in R (v 4.3.0) (*34*). Nuclei with fewer than 800 features or greater than 6000 features were removed to eliminate low quality nuclei and multiplets. In addition, nuclei which had greater than 5% mitochondrial genes were removed. Next, we performed dimensionality reduction by principal component analysis. We utilized the top 30 principal components to calculate the shared nearest neighbor at 0.08 resolution, followed by UMAP dimensionality reduction and visualization. Samples from the mPFC and cerebellum were clustered independently. Each sample retained its sample identity and social status. Any clusters that did not appear in a minimum of 75% of samples or in which 75% or more of the cells belonged to a single animal were removed. Clusters were assigned to known cell types using known cell type specific marker genes.

### Identification of Region-Specific panels of Immediate Early Genes

We started by examining 139 genes previously identified as immediate early genes (*14*). We identified cells which contained at least one copy of a given gene and considered a given IEG socially-responsive if the number of identified neurons increased within any cluster by at least two-fold (with one percent added to denominator) following social interaction (relative to control mice) within pooled experimental groups. Most clusters of cells demonstrated a social interaction-dependent increase of some, but not all IEGs. Because we observed marked region-specific differences in socially responsive IEGs we utilized region-specific panels which included IEGs which displayed a greater than 2-fold increase in expression following social interaction. IEG+ neurons in the mPFC were defined as cells expressing four or more socially-responsive IEGs (43 genes) at above the 95^th^ percentile control level, and IEG+ neurons in the cerebellum were defined as those neurons which expressed two or more socially-responsive IEGs (18 genes) at above the 95^th^ percentile.

### Comparison of IEG expression patterns by region and cell type

To compare the pattern of IEG expression changes driven by social interaction we then determined the fold-increase in the proportion of IEG+ neurons within each identified population of neurons for socially-responsive IEG (using 51 IEGs found to be socially-responsive in either the mPFC or cerebellum). As some smaller clusters had 0 IEG+ cells in the control condition we conservatively added 1 percent to the numerator and denominator. We averaged the fold-difference calculated for each social mouse compared to every control mouse, then calculated and plotted the average across social mice for each condition as a heatmap of neuron types (y) and IEGs (x) sorted by average fold-increase observed in the mPFC. We generated similarity matrices corresponding to Pearson’s pairwise correlations calculated between vectors containing fold-increases in expression (social/control) of each IEG for each mPFC and cerebellar cell type at 10 and 35 minutes which was then averaged across samples.

To further delineate cell-type or regional differences in socially-responsive IEGs we generated counts matrices of socially-responsive IEGs pooled by excitatory or inhibitory neurons from each animal, to generate a matrix corresponding to the expression of all socially-responsive IEGs (51 in total) across all control, 10 minute post-social, and 35 minute post-social animals. We next performed dimensionality reduction via PCA analysis. We found that the first three principal components explained over 80% of the variance. Visualization of the position of each sample in 3-dimensional component space revealed that samples clustered by region and cell-type. To examine how social interaction alters IEG expression we separated each group of samples by region and cell-type (excitatory or inhibitory) and normalized their position by subtracting the mean control position from each sample to visualize the trajectory of social interaction-driven IEG expression. We then compared the trajectories between mice both within and across condition (cell-type and region) by calculating the cosine similarity between each pair of vectors.

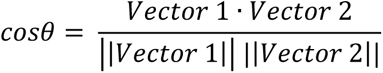

### Quantification of a distributed social ensemble

We quantified the composition of social ensembles at 10 and 35 minutes by determining the excess proportion of IEG+ neurons in each cell type by dividing the proportion of IEG+ neurons in the control condition by the proportion of neurons identified in the same cluster following social interaction, then normalizing to the number of cells identified assigned to the cluster in the social condition.

### Statistics

Differences between social and control group IEG positive proportions were determined by comparing the real proportional increase between experimental and control groups to a distribution of shuffled values for each cell type. For shuffling, the number of real cells belonging to each experimental group were selected from within each cell type and the proportion of IEG+ cells in each randomly selected cell population were calculated. The same comparisons were then calculated for the shuffled data as was done in real data. This was repeated 1000 times to create a distribution of 1000 shuffled values for each cell type. To compare patterns of IEG expression across cell types, we calculated Pearson’s pairwise correlation coefficients and the cosine similarity as described, then performed one-way ANOVA with multiple comparisons to calculate differences between groups using GraphPad Prism (v 9.4.1). Two-way ANOVAs were also calculated in GraphPad Prism.

## Acknowledgements

This work was supported by National Institutes of Health (5K08NS105938 to N.A.F.) and SFARI Bridge to Independence award (to N.A.F.) and a generous donation from Edward Elliott.

---

**Supp. Figure 1:**
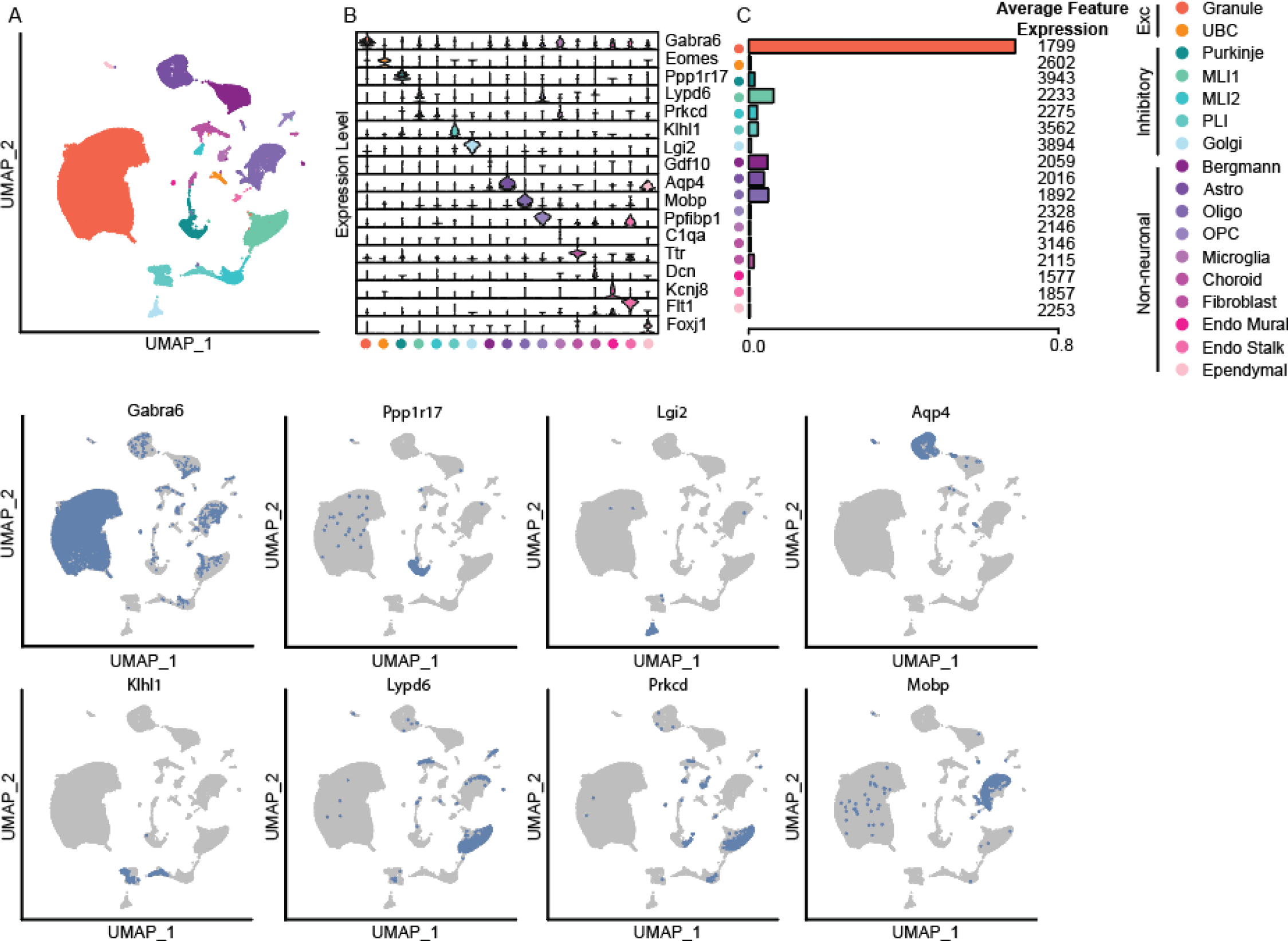
Identification of heterogeneous cerebellar neurons. A) UMAP visualization of clustered cerebellar cells based on nearest neighbor calculations of 90,719 single-nuclei transcriptomes from WT mice (3 early social, 6 late social; 5 not-social). Cell types were annotated based on marker-gene expression. B) Violin plot showing the expression level of known marker genes across cell types (*17*). C) Bar plot of the proportion of cells belonging to each cell type with text representing the average number of genes expressed by cells of that type (Granule – 68.8%, UBC – 0.49%, Purkinje – 1.5%, MLI1 – 6.4%, MLI2 – 2.2%, PLI – 2.4%, Golgi – 0.6%, Astro – 4.0%, Oligo – 5.1%, OPC – 0.5%, Microglia – 0.4%, Bergmann – 4.9%, Choroid – 0.5%, Fibroblast – 1.3%, Endo Mural – 0.2%, Endo Stalk – 0.3%, Ependymal – 0.3%). D) UMAP visualization as in A, with cells colored based on high level of expression (> ∼60% max) of cell-type specific marker genes.

**Supp. Figure 2:**
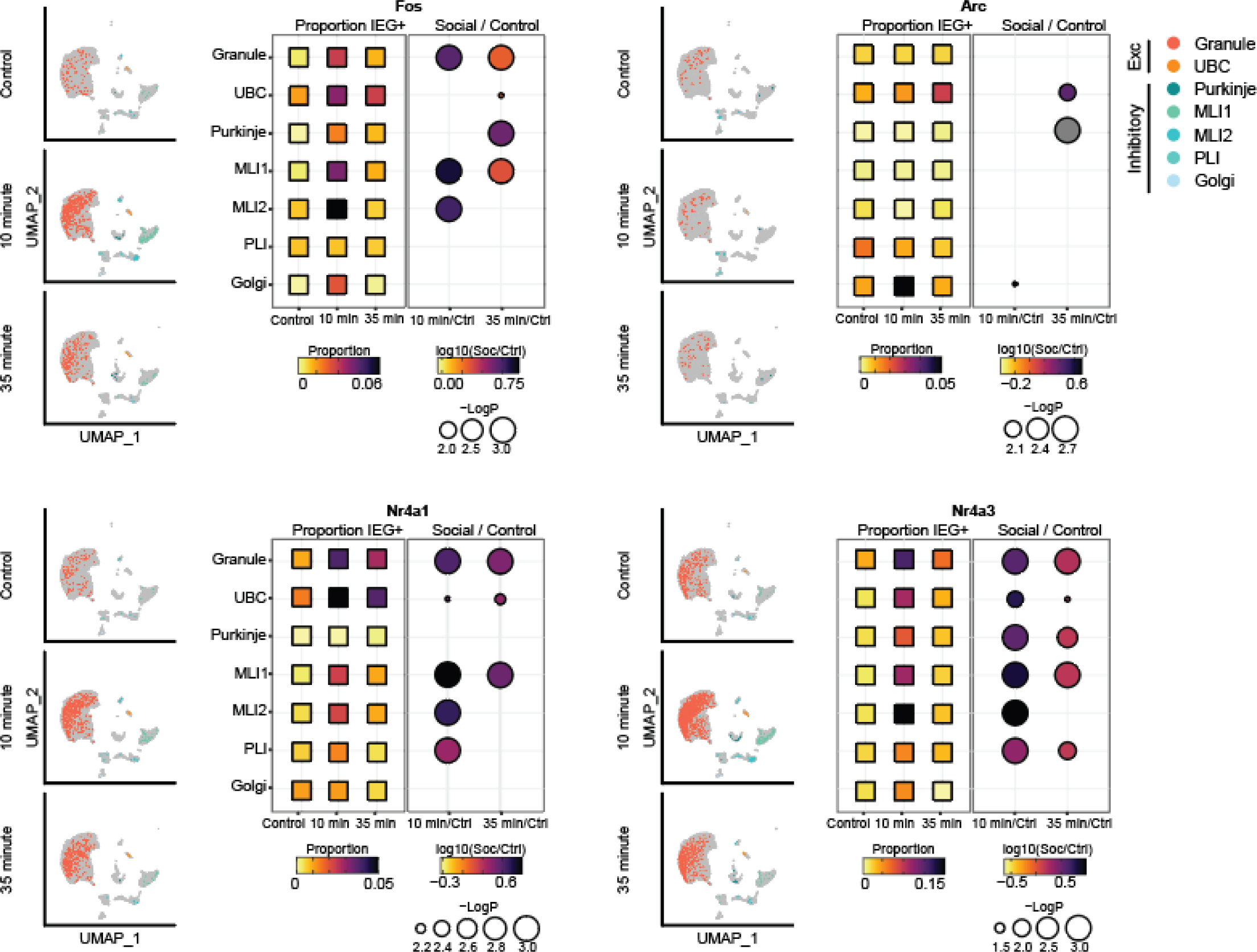
Cerebellar IEG expression increases in heterogeneous cells following social interaction. Cells expressing Fos, Arc, Nr4a1, and Nr4a3 (> 95^th^ percentile) are plotted for each timepoint and quantified. Individual colored dots represent individual neurons expressing corresponding IEG, overlaid on clustered dataset randomly down sampled to show equivalent cell counts in each condition. Corresponding dot plots represent the proportion of cells expressing the corresponding IEG at 10 minutes and 35 minutes post social interaction, and the ratio of the proportions of positive cells in the post-social vs control conditions (also see Supp. Table 1; p-values calculated based on the shuffled data).

**Supp. Figure 3:**
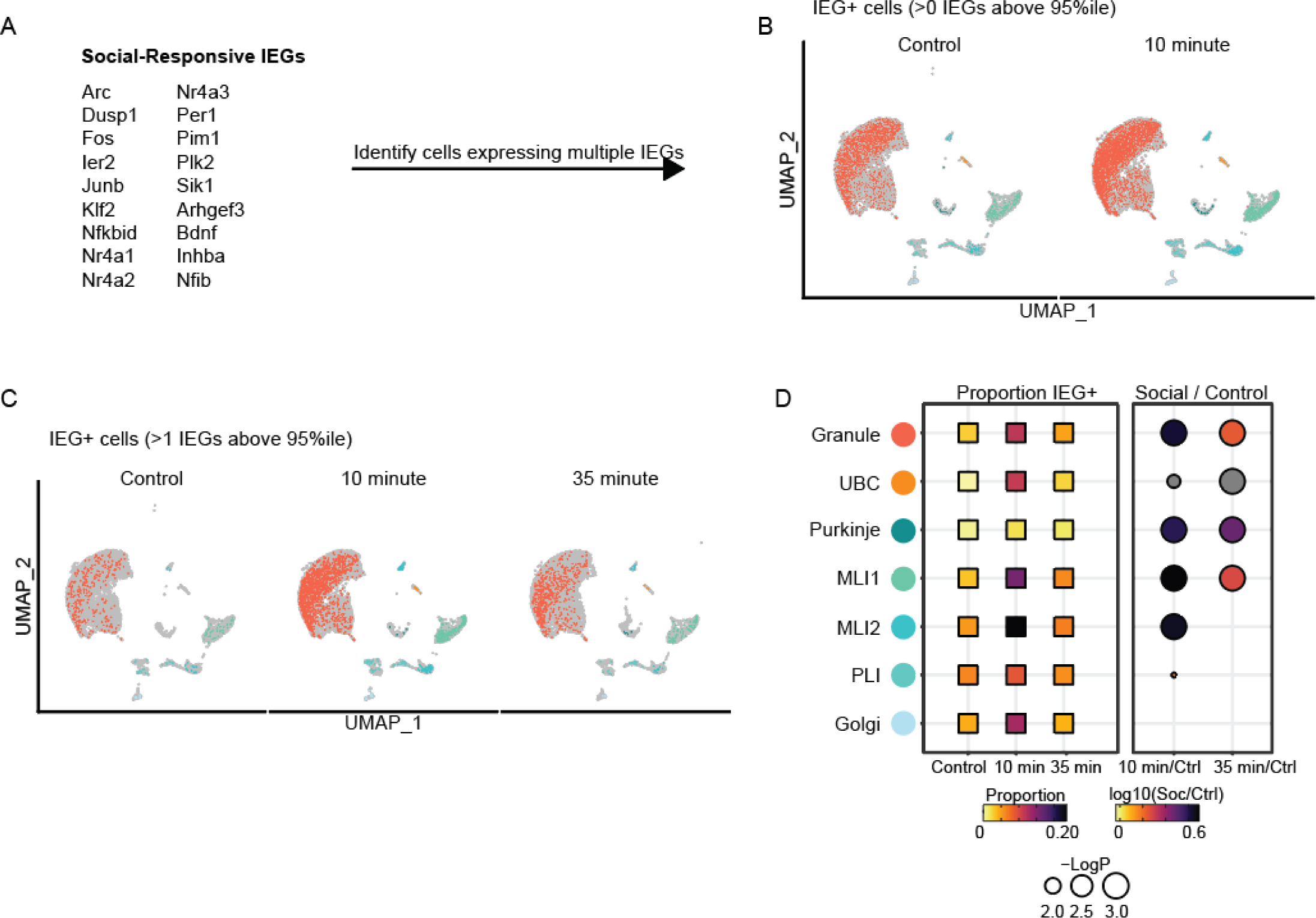
Recruitment of heterogeneous cerebellar neurons following social interaction. We utilized a panel of 18 socially-responsive IEGs in the mPFC (>2-fold expression in social condition compared to control). B) UMAPs showing the proportion of cells of each type which express at least a single IEG (top) or 2 or more IEGs (bottom) in the control and 10-minute post-social condition. C) Proportion of cells expressing 4 or more IEGs above the 95th percentile of expression in the control group. UMAPs are randomly down sampled by condition to show the same number of cells for each condition (B&C). D) Boxes on left are colored to represent the proportion of IEG+ neurons in each cell type in control or social conditions (n = 34,710 neurons from 5 control mice, 27,468 neurons from 6 35-minute post social mice and 12,699 neurons from 3 ten-minute post social mice). We then calculated the proportion of IEG+ neurons at each post-social timepoint. Circles are colored to represent the ratio of IEG+ cells in each social condition relative to the control condition. The diameter of each circle represents Log10 p-value (also see Supp. Table 4; p-values calculated based on shuffled data).

**Supp. Figure 4:**
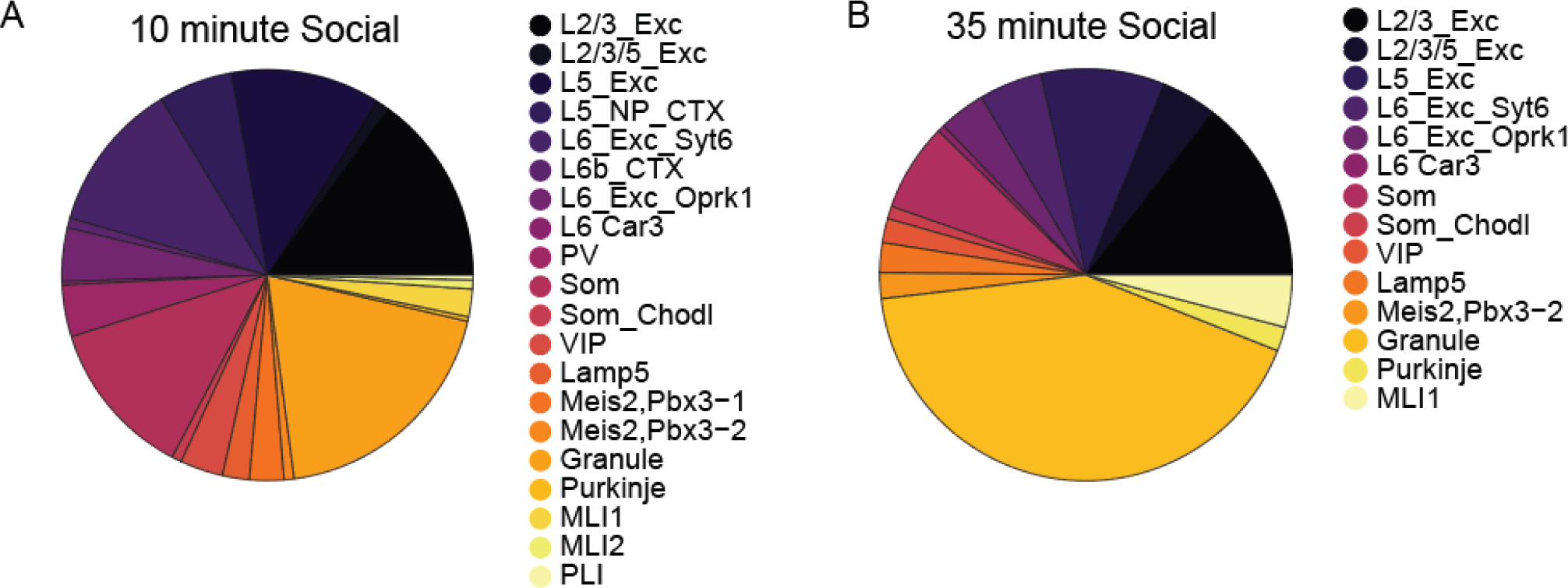
A distributed social ensemble. We quantified the relative contribution of each identified neuronal subtype from the mPFC or cerebellum to the representation of social information (see methods for calculation). We included only clusters in which the proportion of IEG+ neurons increased significantly following social interaction. A) Contribution of each cell type to the overall increase in IEG+ neurons following social interaction at 10 minutes (L2/3 Exc – 14.9%, L2/3/5 Exc – 1.09%, L5 Exc – 11.7%, L5 NP CTX 5.82%, L6 Exc Syt6 – 11.8%, L6b CTX – 0.78%, L6 Exc Oprk1 – 4.19%, L6 Car3 – 0.27%, PV - 4.06%, Som – 12.4%, Som Chodl – 0.76%, VIP – 3.38%, Lamp5 – 2.11%, Meis2, Pbx3-1 – 2.69%, Meis2, Pbx3-2 – 0.79%, Granule – 19.3%, Purkinje – 0.35%, MLI1 – 2.17%, MLI2 – 0.70%, PLI – 0.37%). B) Contribution of each cell type to the overall increase in IEG+ neurons following social interaction at 35 minutes (L2/3 Exc – 14.4%, L2/3/5 Exc – 4.36%, L5 Exc - 9.69%, L6 Exc Syt6 – 4.93%, L6 Exc Oprk1 – 3.65%, L6 Car3 – 0.57%, Som – 6.91%, Som Chodl – 0.99%, VIP – 1.89%, Lamp5 – 2.33%, Meis2,Pbx3-2 – 2.03%, Granule – 42.1%, Purkinje – 1.89%, MLI1 – 4.10%).

**Supp. Table 1.**

| Comparison | Cell type | Fos |  | Arc |  | Nr4a1 |  | Nr4a3 |  |
| --- | --- | --- | --- | --- | --- | --- | --- | --- | --- |
|  |  | ratio | p-value | ratio | p-value | ratio | p-value | ratio | p-value |
| 10 min. /Ctrl | Meis2,Pbx3-2 | 3.95 | 0.001 | 7.00 | 0.001 | 5.39 | 0.001 | 1.54 | 0.063 |
| 10 min. /Ctrl | Meis2,Pbx3-1 | 3.17 | 0.001 | 2.84 | 0.001 | 3.38 | 0.001 | 5.56 | 0.001 |
| 10 min. /Ctrl | Lamp5 | 4.16 | 0.001 | Inf | 0.001 | 2.92 | 0.003 | 3.47 | 0.001 |
| 10 min. /Ctrl | VIP | 5.28 | 0.001 | 8.67 | 0.001 | 3.96 | 0.001 | 10.73 | 0.001 |
| 10 min. /Ctrl | Som_Chodl | 5.70 | 0.001 | 8.54 | 0.001 | 21.36 | 0.001 | 1.42 | 0.013 |
| 10 min. /Ctrl | Som | 7.12 | 0.001 | 7.54 | 0.001 | 5.65 | 0.001 | Inf | 0.001 |
| 10 min. /Ctrl | PV | 7.20 | 0.001 | 6.66 | 0.001 | 2.85 | 0.001 | 6.98 | 0.001 |
| 10 min. /Ctrl | L6 Car3 | 5.13 | 0.001 | 11.55 | 0.001 | Inf | 0.001 | 10.27 | 0.001 |
| 10 min. /Ctrl | L6_Exc_Oprk1 | 4.33 | 0.001 | 7.09 | 0.001 | 5.88 | 0.001 | 3.16 | 0.001 |
| 10 min. /Ctrl | L6b_CTX | 3.68 | 0.001 | 8.11 | 0.001 | 3.32 | 0.001 | 2.46 | 0.001 |
| 10 min. /Ctrl | L6_Exc_Syt6 | 5.65 | 0.001 | 9.82 | 0.001 | 3.69 | 0.001 | 3.27 | 0.001 |
| 10 min. /Ctrl | L5_NP_CTX | 9.12 | 0.001 | 14.92 | 0.001 | 2.98 | 0.001 | 2.09 | 0.004 |
| 10 min. /Ctrl | L5_Exc | 5.78 | 0.001 | 10.29 | 0.001 | 3.67 | 0.001 | 4.43 | 0.001 |
| 10 min. /Ctrl | L2/3/5_Exc | 5.38 | 0.001 | 34.21 | 0.001 | 2.68 | 0.001 | 4.68 | 0.001 |
| 10 min. /Ctrl | L2/3_Exc | 5.47 | 0.001 | 8.92 | 0.001 | 4.21 | 0.001 | 4.65 | 0.001 |
| 35 min. / Ctrl | Meis2,Pbx3-2 | 1.45 | 0.046 | 3.86 | 0.001 | 5.37 | 0.001 | 0.38 | 1 |
| 35 min. / Ctrl | Meis2,Pbx3-1 | 1.08 | 0.287 | 0.78 | 0.879 | 1.22 | 0.181 | 1.75 | 0.001 |
| 35 min. / Ctrl | Lamp5 | 2.10 | 0.006 | Inf | 0.001 | 0.88 | 0.717 | 1.47 | 0.046 |
| 35 min. / Ctrl | VIP | 1.88 | 0.051 | 0.78 | 0.801 | 0.63 | 0.913 | 1.04 | 0.392 |
| 35 min. / Ctrl | Som_Chodl | 3.74 | 0.001 | 1.07 | 0.21 | 1.60 | 0.004 | 1.60 | 0.003 |
| 35 min. / Ctrl | Som | 1.02 | 0.453 | 2.61 | 0.001 | 0.00 | 1 | Inf | 0.001 |
| 35 min. / Ctrl | PV | 2.00 | 0.001 | 1.40 | 0.231 | 2.10 | 0.001 | 0.93 | 0.705 |
| 35 min. / Ctrl | L6 Car3 | 1.70 | 0.042 | 3.51 | 0.004 | NaN | 1 | 2.56 | 0.013 |
| 35 min. / Ctrl | L6_Exc_Oprk1 | 1.54 | 0.001 | 2.29 | 0.001 | 1.76 | 0.001 | 1.35 | 0.005 |
| 35 min. / Ctrl | L6b_CTX | 1.26 | 0.061 | 1.90 | 0.001 | 1.58 | 0.001 | 1.21 | 0.085 |
| 35 min. / Ctrl | L6_Exc_Syt6 | 1.97 | 0.001 | 2.05 | 0.001 | 1.31 | 0.06 | 1.18 | 0.211 |
| 35 min. / Ctrl | L5_NP_CTX | 2.19 | 0.002 | 3.94 | 0.001 | 1.18 | 0.127 | 0.97 | 0.631 |
| 35 min. / Ctrl | L5_Exc | 2.14 | 0.001 | 2.35 | 0.001 | 1.67 | 0.001 | 1.68 | 0.001 |
| 35 min. / Ctrl | L2/3/5_Exc | 2.39 | 0.001 | 7.21 | 0.001 | 1.66 | 0.001 | 2.52 | 0.001 |
| 35 min. / Ctrl | L2/3_Exc | 2.12 | 0.001 | 2.31 | 0.001 | 1.50 | 0.001 | 1.84 | 0.001 |

Note: Values represent social/control for each cell type for individual IEGs and p-values in the mPFC (see Figure 4).

**Supp. Table 2.**

| Comparison | Cell type | Fold-change | log10(Fold-change) | p-value |
| --- | --- | --- | --- | --- |
| 10 min. / Ctrl | L2/3_Exc | 8.80 | 0.94 | 0.001 |
| 10 min. / Ctrl | L2/3/5_Exc | 9.34 | 0.97 | 0.001 |
| 10 min. / Ctrl | L5_Exc | 8.27 | 0.92 | 0.001 |
| 10 min. / Ctrl | L5_NP_CTX | 19.89 | 1.30 | 0.001 |
| 10 min. / Ctrl | L6_Exc_Syt6 | 10.65 | 1.03 | 0.001 |
| 10 min. / Ctrl | L6b_CTX | 4.91 | 0.69 | 0.001 |
| 10 min. / Ctrl | L6_Exc_Oprk1 | 7.11 | 0.85 | 0.001 |
| 10 min. / Ctrl | L6 Car3 | 5.60 | 0.75 | 0.001 |
| 10 min. / Ctrl | PV | 9.13 | 0.96 | 0.001 |
| 10 min. / Ctrl | Som | 48.98 | 1.69 | 0.001 |
| 10 min. / Ctrl | Som_Chodl | 11.75 | 1.07 | 0.012 |
| 10 min. / Ctrl | VIP | 18.57 | 1.27 | 0.001 |
| 10 min. / Ctrl | Lamp5 | 13.53 | 1.13 | 0.001 |
| 10 min. / Ctrl | Meis2,Pbx3-1 | 6.07 | 0.78 | 0.001 |
| 10 min. / Ctrl | Meis2,Pbx3-2 | 6.03 | 0.78 | 0.001 |
| 35 min. / Ctrl | L2/3_Exc | 2.28 | 0.36 | 0.001 |
| 35 min. / Ctrl | L2/3/5_Exc | 3.39 | 0.53 | 0.001 |
| 35 min. / Ctrl | L5_Exc | 2.18 | 0.34 | 0.001 |
| 35 min. / Ctrl | L5_NP_CTX | 0.88 | -0.06 | 0.727 |
| 35 min. / Ctrl | L6_Exc_Syt6 | 1.45 | 0.16 | 0.004 |
| 35 min. / Ctrl | L6b_CTX | 1.33 | 0.13 | 0.205 |
| 35 min. / Ctrl | L6_Exc_Oprk1 | 2.06 | 0.31 | 0.001 |
| 35 min. / Ctrl | L6 Car3 | 2.38 | 0.38 | 0.015 |
| 35 min. / Ctrl | PV | 1.40 | 0.15 | 0.084 |
| 35 min. / Ctrl | Som | 7.84 | 0.89 | 0.001 |
| 35 min. / Ctrl | Som_Chodl | 2.40 | 0.38 | 0.001 |
| 35 min. / Ctrl | VIP | 2.74 | 0.44 | 0.001 |
| 35 min. / Ctrl | Lamp5 | 4.42 | 0.65 | 0.001 |
| 35 min. / Ctrl | Meis2,Pbx3-1 | 1.05 | 0.02 | 0.451 |
| 35 min. / Ctrl | Meis2,Pbx3-2 | 2.01 | 0.30 | 0.012 |

Note: Values represent those of social /control, as displayed in Figure 5D, for the cell type specific increases in IEG panel expression.

**Supp. Table 3.**

| Comparison | Cell type | Fos |  | Arc |  | Nr4a1 |  | Nr4a3 |  |
| --- | --- | --- | --- | --- | --- | --- | --- | --- | --- |
|  |  | ratio | p-value | ratio | p-value | ratio | p-value | ratio | p-value |
| 10 min. / Ctrl | Golgi | 9.10 | 0.418 | 5.05 | 0.014 | 1.01 | 0.589 | 2.53 | 0.088 |
| 10 min. / Ctrl | PLI | 1.01 | 0.534 | 0.60 | 0.977 | 2.42 | 0.001 | 2.27 | 0.001 |
| 10 min. / Ctrl | MLI2 | 5.78 | 0.001 | 0.00 | 1 | 4.95 | 0.001 | 9.77 | 0.001 |
| 10 min. / Ctrl | MLI1 | 7.27 | 0.001 | 0.55 | 0.99 | 7.88 | 0.001 | 6.68 | 0.001 |
| 10 min. / Ctrl | Purkinje | 7.26 | 0.056 | NaN | 1 | NaN | 1 | 3.57 | 0.001 |
| 10 min. / Ctrl | UBC | 2.58 | 0.11 | 1.29 | 0.114 | 3.43 | 0.007 | 5.79 | 0.013 |
| 10 min. / Ctrl | Granule | 5.07 | 0.001 | 0.95 | 0.836 | 4.11 | 0.001 | 3.79 | 0.001 |
| 35 min. / Ctrl | Golgi | 1.38 | 0.556 | 0.92 | 0.646 | 0.46 | 0.979 | 0.23 | 0.997 |
| 35 min. / Ctrl | PLI | 0.92 | 0.656 | 0.41 | 0.995 | 0.73 | 0.967 | 1.44 | 0.01 |
| 35 min. / Ctrl | MLI2 | 0.89 | 0.817 | 0.89 | 0.793 | 2.07 | 0.101 | 1.58 | 0.125 |
| 35 min. / Ctrl | MLI1 | 2.28 | 0.001 | 0.80 | 0.982 | 3.35 | 0.001 | 1.51 | 0.001 |
| 35 min. / Ctrl | Purkinje | 4.51 | 0.001 | Inf | 0.001 | Inf | 0.001 | 1.46 | 0.005 |
| 35 min. / Ctrl | UBC | 1.94 | 0.03 | 2.59 | 0.006 | 2.59 | 0.006 | 1.94 | 0.039 |
| 35 min. / Ctrl | Granule | 2.07 | 0.001 | 0.98 | 0.729 | 2.93 | 0.001 | 1.65 | 0.001 |

Note: Values represent social/control for each cell type for individual IEGs and p-values in the cerebellum (see Supp. figure 3).

**Supp. Table 4.**

| Comparison | Cell type | Fold-change | log10(Fold-change) | p-value |
| --- | --- | --- | --- | --- |
| 10 min. / Ctrl | Granule | 4.21 | 0.62 | 0.001 |
| 10 min. / Ctrl | UBC | Inf | NaN | 0.014 |
| 10 min. / Ctrl | Purkinje | 3.87 | 0.59 | 0.001 |
| 10 min. / Ctrl | MLI1 | 4.90 | 0.69 | 0.001 |
| 10 min. / Ctrl | MLI2 | 4.43 | 0.65 | 0.001 |
| 10 min. / Ctrl | PLI | 1.46 | 0.16 | 0.024 |
| 10 min. / Ctrl | Golgi | 3.03 | 0.48 | 0.074 |
| 35 min. / Ctrl | Granule | 1.73 | 0.24 | 0.001 |
| 35 min. / Ctrl | UBC | Inf | NaN | 0.001 |
| 35 min. / Ctrl | Purkinje | 3.01 | 0.48 | 0.001 |
| 35 min. / Ctrl | MLI1 | 1.88 | 0.27 | 0.001 |
| 35 min. / Ctrl | MLI2 | 1.26 | 0.10 | 0.069 |
| 35 min. / Ctrl | PLI | 0.93 | -0.03 | 0.689 |
| 35 min. / Ctrl | Golgi | 0.92 | -0.04 | 0.705 |

Note: Values represent those of social /control, as displayed in Supp. figure 4D, for the cell type specific increases in IEG panel expression.

